# Critical role of Cas9 aggregation on *in vitro* DNA cleavage

**DOI:** 10.64898/2025.12.23.696176

**Authors:** Diego Cora, Ana Seijas, Wajih Al-Soufi, Laura Sánchez, Álvaro J. Arana, Mercedes Novo

## Abstract

The CRISPR/Cas9 system is a powerful genome-editing tool widely used in molecular biology and gene therapy, whose efficiency strongly depends on the physicochemical properties of the Cas9 ribonucleoprotein complex. Optimizing Cas9 activity remains essential for reliable genome- editing applications, yet the factors limiting its *in vitro* cleavage efficiency are not fully understood. Among these, protein aggregation has been suggested to critically impair Cas9 functionality, although its role has not been systematically analysed.

Here, we investigate Cas9 aggregation under different environmental conditions and evaluate its impact on *in vitro* DNA cleavage efficiency. Using fluorescently labelled Cas9 and single-molecule fluorescence techniques, we quantify aggregation as a function of buffer composition, ionic strength, salt concentration and sgRNA presence, and relate these properties to cleavage activity. Our results show that Cas9 aggregation significantly reduces DNA cleavage efficiency, with higher aggregation levels consistently correlating with lower activity. In contrast, buffers with higher ionic strength or stabilizing components reduce aggregation and enhance Cas9 performance.

Overall, this study demonstrates that Cas9 aggregation plays a critical role in determining *in vitro* cleavage efficiency and highlights the importance of controlling protein aggregation to optimize CRISPR/Cas9-based genome-editing applications and delivery strategies.

**GRAPHICAL ABSTRACT:** 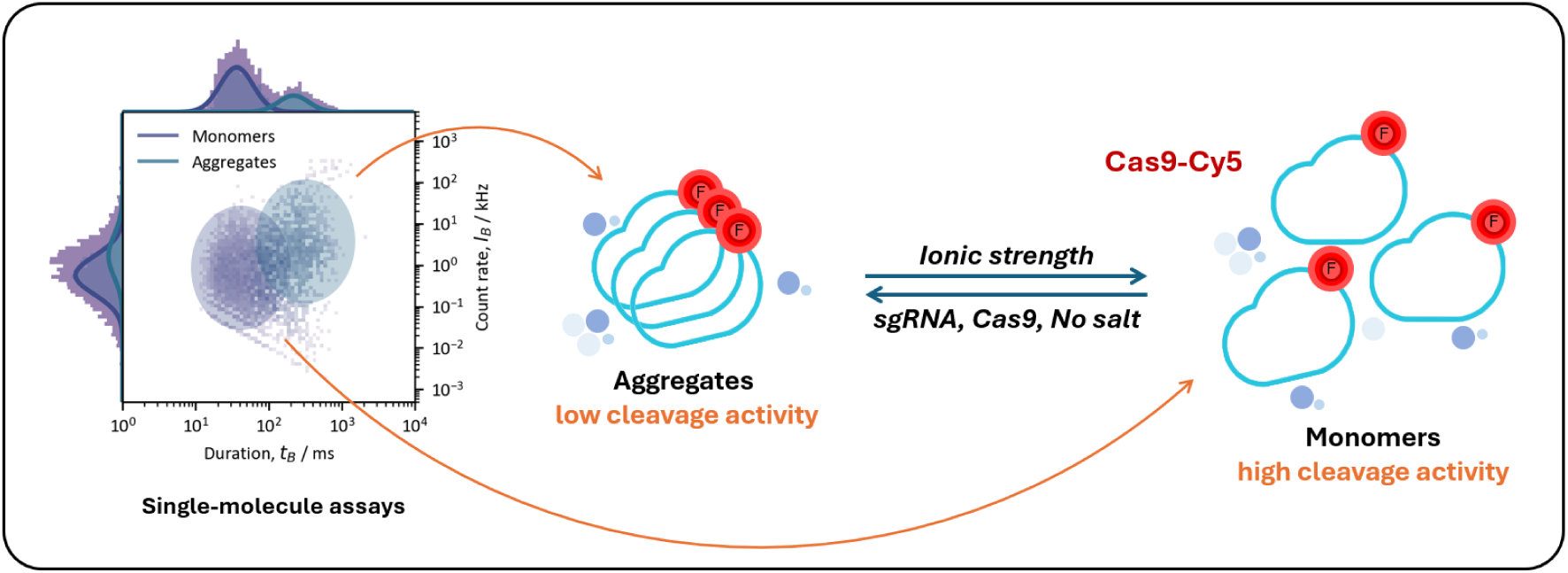

## 1. INTRODUCTION

CRISPR/Cas9 (Clustered Regularly Interspaced Short Palindromic Repeats/CRISPR-associated protein 9) is a well-known genome-editing tool derived from the adaptive immune system of bacteria (1, 2). Cas9, a large multi-domain enzyme, recognizes the protospacer adjacent motif (PAM), unwinds the DNA, and facilitates site-specific cleavage. This cleavage triggers cellular repair mechanisms, enabling precise gene editing (3). This system employs a single guide RNA (sgRNA) to form a ribonucleoprotein (RNP) complex with the Cas9 endonuclease, directing it to a complementary DNA sequence where the double-stranded break is introduced (4). Owing to its high efficiency and programmability, CRISPR/Cas9 has become indispensable in fields ranging from basic biology to clinical therapeutics.

Despite the broad application of CRISPR/Cas9, relatively little attention has been given to the physicochemical properties of the Cas9 protein itself, particularly regarding its tendency to aggregate under various environmental conditions (5). Aggregation can significantly compromise Cas9’s efficiency in gene editing, by reducing the availability of functional monomeric Cas9 and hindering its encapsulation into delivery vehicles or cellular uptake due to increased particle size (5). Although scattered reports have described aggregation-related behaviour of Cas9, a systematic understanding of this phenomenon is still lacking. For example, Manzano *et al*. (6) described shear-induced aggregation near the isoelectric point of Cas9, while Nguyen *et al*. (7) reported significant size increase upon sgRNA addition, indicating aggregate formation. Jain *et al*. (8) further demonstrated that aggregation propensity increases near the protein’s isoelectric point and hypothesised potentially impairing sgRNA complexation.

During preliminary studies in our laboratory, focused on delivery systems for CRISPR/Cas9 complexes, we consistently observed unexpectedly high levels of protein aggregation. We also observed a reduction in the *in vivo* genome-editing efficiency across different RNP delivery systems, related with changes in the aggregation state of Cas9 depending on the formulation medium (9). These observations suggested an inverse relationship between Cas9 aggregation and genome-editing efficiency. Together, these findings motivated us to systematically investigate the *in vitro* editing efficiency of Cas9 under standard experimental conditions and to correlate it with the aggregation behaviour of the protein.

Building on these insights, the present study aims to systematically analyse the *in vitro* DNA cleavage efficiency of Cas9 and to relate it to the aggregation state of the protein under different experimental conditions. To this end, agarose gel electrophoresis and capillary electrophoresis were used to quantify Cas9’s cleavage ability under different experimental conditions, while fluorescently labelled Cas9 with Cy5 was used and single-molecule fluorescence experiments were conducted to extract the aggregation information. In our group, we have prior experience applying Fluorescence Correlation Spectroscopy (FCS) to the study of amyloid aggregation processes (10, 11). In this case, we build upon this approach, taking advantage of single-molecule detection (SMD) to separate different populations (monomers and aggregates) and analyse them independently, improving our results. In parallel, our group has extensive experience in genetic editing assays and *in vivo* CRISPR/Cas9 applications (9).

## 2. MATERIALS AND METHODS

### 2.1. Materials

Recombinant SpCas9 proteins were produced and purified by the Proteomics Service of the Andalusian Centre for Developmental Biology (CABD, a joint institute of Pablo de Olavide University and the Spanish National Research Council, Seville, Spain). Two preparations were used in this study: an unlabelled SpCas9 (hereafter referred to as Cas9) and a Cy5-labelled SpCas9 (Cas9*). Both were supplied as stock solutions in 50% (v/v) glycerol/water at concentrations of 1.5 µg/µL for Cas9 and 9 µg/µL for Cas9*.

Single-guide RNAs (sgRNAs) were prepared and purified as described for the DNA cleavage assays. The same sgRNAs used for the *in vitro* DNA cleavage experiments were employed in all SMD/FCS measurements.

Buffers for *in vitro* cleavage experiments were prepared using NaCl, MgCl_2_, KCl, HCl, Tris (tris(hydroxymethyl)aminomethane), HEPES (4-(2-hydroxyethyl)-1-piperazineethanesulfonic acid), BSA (Bovine serum albumin), EDTA (Ethylenediamine tetraacetic acid), DTT (Dithiothreitol) and glycerol. All these reagents were purchased from Merck. To see buffer composition and preparation go to section 2.2.3. The same NaCl and MgCl_2_ were used to prepare solutions for the aggregation studies (to see further explanation go to section 2.3.1).

Phosphate buffered saline (PBS, pH 7.4, 12 mM phosphates, 137 mM NaCl, 3 mM KCl and ionic strength of *I* = 0.18 M) and Tris buffer (10 mM, pH 7.4) were also prepared to be used for sample preparation in the Cas9’s aggregation experiments (each series will indicate which buffer was used). All aqueous solutions were prepared using Milli-Q water.

### 2.2. DNA cleavage assays

#### 2.2.1. DNA extraction, PCR and DNA purification

Firstly, DNA from zebrafish embryos was extracted using Chelex® 100 Resin (Bio-Rad). Individual embryos were resuspended in 120 µL of 10 % Chelex solution in Milli-Q water and incubated at 100 °C for 15 min. Then, 3.1 µL of Proteinase K was added, followed by incubation at 56 °C for 1 h, and a final step at 100 °C for 15 min to inactivate the enzyme.

Primer pairs for the zebrafish *ppifb* gene were designed using NCBI Primer-BLAST (https://www.ncbi.nlm.nih.gov/tools/primer-blast/) based on the reference sequence of *D. labrax.* One amplicon of 459 bp was selected to ensure robust amplification and facilitate sequencing across the target region. Primer selection prioritized specificity, optimal melting temperature (*T*_m_), GC content, and low self-complementarity. The sequences and thermodynamic parameters of the primers are detailed in the Supplementary Information (Table S1).

Polymerase chain reaction (PCR) was performed to amplify the DNA region of interest. For the amplification of the zebrafish *ppifb* gene, reagents were mixed as follows: 10.18 µL of Milli-Q® water, 0.12 µL of both primer forward (*ppifb*-F; 5ʹ 6-FAM–labelled for fragment analysis) and reverse (*ppifb*-R), 0.75 µL of dNTPs (2 µM each), 1.5 µL of 10X PCR Gold Buffer (Applied Biosystems), 1.3 µL of MgCl_2_ solution 25 mM (Applied Biosystems), 0.2 µL of AmpliTaq Gold™ DNA Polymerase 1000 U (Applied Biosystems) and 0.8 µL of previously extracted DNA. PCR tubes were then introduced in a thermocycler, and the following program was set: samples were maintained at 25 °C for 10 min and then heated to 95 °C for 10 min, followed by 35 cycles of 94 °C for 45 s, 62 °C for 50 s and 72 °C for 50 s, and a final step of 72 °C for 10 min was performed.

Two DNA samples corresponding to the same *ppifb* amplicon were used as substrates: one derived from a wild-type individual (*ppifb* WT) and the other from a heterozygous edited allele (*ppifb*-het). The *ppifb*-het (hereafter referred to as *ppifb*) template used in the *in vitro* assays was derived from a heterozygous offspring obtained by crossing an adult CRISPR/Cas9-edited F0 individual with a wild-type fish. The *ppifb* amplicon was included to introduce within-sample sequence heterogeneity and to evaluate the robustness of the *in vitro* cleavage readout in the presence of minor allelic variation within the amplified region, including variants located outside the guide-binding regions and, for one sgRNA, within the protospacer sequence.

Efficient PCR was confirmed by performing agarose gel (1 %) electrophoresis. DNA was then purified using MinElute® PCR Purification Kit from QIAGEN according to the manufacturer’s instructions. Finally, DNA concentration was measured in a NanoDrop® ND-1000 Spectrophotometer and stored at −20 °C until usage.

#### 2.2.2. sgRNA design, synthesis and preparation

sgRNAs targeting the zebrafish *ppifb* locus were designed in silico using the CRISPRscan online tool and the CRISPOR web server, selecting target sites of the form N₂₀–NGG within the coding region. Candidate guides were ranked according to their predicted on-target efficiency (CRISPRscan score), GC content and position within the gene, together with CRISPOR-predicted specificity metrics (MIT and CFD scores, number of off-targets) and Doench’16 activity scores. Guides with the lowest predicted off-target activity were prioritized. Three sgRNAs with the highest predicted activity and suitable sequence features were selected for experimental validation, and the corresponding oligonucleotides were synthesized for *in vitro* transcription (**Table 1** & **Table 2**).

**Table 1.**
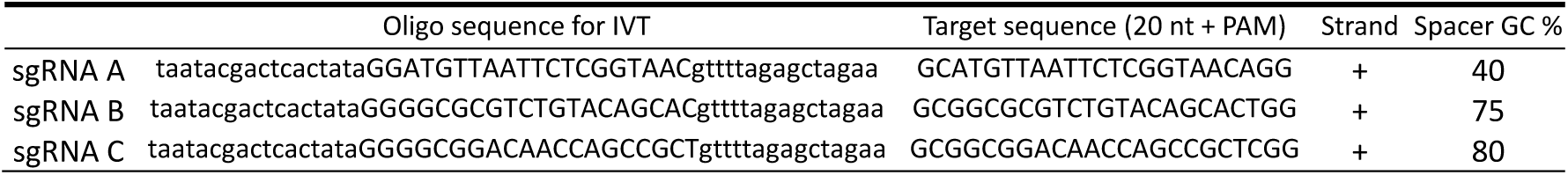
Sequence features of the sgRNAs targeting zebrafish *ppifb*.

**Table 2.** CRISPRscan- and CRISPOR-predicted activity and specificity scores for the sgRNAs.

|  | CRISPRscan<br>score | MIT specificity<br>score | CFD specificity<br>score | Doench'16<br>predicted efficiency | Off-targets<br>(0–1–2–3–4 MM)* | Total off-targets |
| --- | --- | --- | --- | --- | --- | --- |
| sgRNA A | 44 | 99 | 99 | 43 | 0–0–0–1–17 | 18 |
| sgRNA B | 62 | 92 | 97 | 40 | 0–0–0–2–18 | 20 |
| sgRNA C | 66 | 98 | 99 | 56 | 0–0–0–1–8 | 9 |

\*Number of predicted off-target sites in the genome with 0, 1, 2, 3 and 4 mismatches (MM) relative to the spacer sequence (as predicted by CRISPOR).

Templates for *in vitro* transcription were generated by PCR amplification mixing 5 µL of iProof™ High-Fidelity PCR buffer (Bio-Rad) 5X, 1.25 µL of dNTPs (2 mM each), 1 µL of sgRNA primer 10 mM, 1 µL of universal primer 10 mM, 0.25 µL of Taq iProof™ High-Fidelity DNA Polymerase (Bio- Rad and 16.5 µL of Milli-Q® water. Samples were then introduced in a thermocycler, and the following program was set: firstly, they were heated to 98 °C for 3 min, followed by 30 cycles of 98 °C for 30 s, 45 °C for 30 s and 72 °C for 20 s, and a final step of 72 °C for 5 min was performed. *In vitro* transcription was performed using the MAXIscript™ T7 Transcription Kit (Thermo Fisher Scientific) with some protocol modifications: a mix containing 3 µL of Nuclease-Free water, 3 µL of PCR product, 1 µL of 10X Transcription Buffer, 1 µL of Enzyme Mix and 0.5 µL of each ATP, GTP, CTP and UTP (10 mM) was made, mixed and stored overnight at 37 °C. Afterwards, 1.5 µL of TURBO DNase (2 U/µL) was added to the samples and they were incubated at 37 °C for 30 min. Precipitation was performed with 2 µL of EDTA 0.5 M, 15 µL of LiCl 5 M and 150 µL of 100 % ethanol at −80 °C for 3 h 45 min. Then, a centrifugation of 30 min at 13000 rpm and 4 °C was performed and a wash step with 200 µL of ethanol followed by centrifugation at 13000 rpm and 4 °C for 20 min were conducted twice. RNA was finally resuspended in Nuclease-Free Water and RNA concentration was measured in a NanoDrop® ND-1000 Spectrophotometer and stored at −80 °C until usage.

#### 2.2.3. In vitro cleavage experiments

Three different buffers described in the literature (12–18) were prepared to study *in vitro* editing efficiency: 10X NEBuffer (NB), containing NaCl 1 M, MgCl_2_ 0.1 M, Tris-HCl 0.5 M, BSA 1 mg/mL with pH 7.9 (ionic strength of *I* = 0.18 M) in Milli-Q; 10X Reaction Buffer (RB), containing NaCl 1 M, MgCl_2_ 0.05 M, HEPES 0.2 M, EDTA 0.001 M (*I* = 0.14 M) in Milli-Q; and 1X Cleavage Buffer (CB), containing KCl 0.1 M, MgCl_2_ 0.005 M, Tris-HCl 0.02 M, 5% Glycerol and DTT 0.001 M with pH 7.5 (*I* = 0.12 M) in Milli-Q water.

The following protocol was performed for the *in vitro* editing digestion obtaining a final volume of 10 µL. Milli-Q water and buffer were mixed to obtain 1X buffer in each case (note that buffer CB is already 1X so no dilution was needed). Synthesized sgRNA and commercial Cas9 enzyme were added in a molar ratio of 1:1 (in our experiments 0.3 μM was the concentration used) and the mixture was incubated for 15 min at room temperature to allow the formation of the RNP complex. DNA was then added (between 100 and 250 ng depending on the experiment), and digestion was performed at 37 °C for 15 min or 1 h. Finally, reaction was stopped by freezing the samples and analysis was performed by the techniques described in section 2.2.4. Control samples with DNA but neither Cas9 or sgRNA were made by the protocol described above and used in agarose gel and capillary electrophoresis but not in data analysis.

#### 2.2.4. Analysis techniques for in vitro DNA cleavage

DNA cleavage analysis was carried out by two methods, using agarose gel (1 %) electrophoresis for a qualitative determination of the genome-editing output, and by capillary electrophoresis fragment analysis for a quantitative determination of the DNA cleavage efficiency.

Agarose gel electrophoresis analysis was performed to observe the results of cleavage assays and to guide the dilution range required for subsequent fragment analysis. To do so, 5μL of the *in vitro* editing digestion product were mixed with bromophenol blue loading dye, loaded into a 1% agarose gel prepared in 0.5× TAE and stained with SYBR™ Safe DNA Gel Stain (Thermo Fisher Scientific), and electrophoresed at 120 V in 0.5× TAE buffer. GeneRuler 100 bp DNA Ladder was loaded in the first column as a reference for size DNA fragments.

In addition, samples obtained by the *in vitro* cleavage assays were diluted 1:20 - 1:50 in Milli-Q water to perform fragment analysis. Diluted samples were then mixed with Hi-Di™ formamide and an internal size standard (GeneScan™; Applied Biosystems) according to the manufacturer’s instructions, denatured (95 °C for 3 min) and immediately cooled on ice prior to loading. Raw fragment electropherograms generated by the ABI 3500xL Genetic Analyzer (Applied Biosystems) and GeneMapper™ Software v4.1 were exported, containing peak size (in base pairs, bp), peak height, and peak area information. These files were processed using a custom Python workflow developed for this study.

Peaks corresponding to the expected cleavage products were extracted using predefined size windows: 175–195 bp (sgRNA A), 220–240 bp (sgRNA B), and 285–305 bp (sgRNA C), as well as the intact DNA peak (440–470 bp). Peak areas within these windows were summed, and cleavage efficiency was calculated as the fraction of total fragment area attributable to the cleavage-derived products.

A structured dataset was built and used to produce summary statistics (mean ± standard deviation) to generate bar plots illustrating the effect of each factor on cleavage performance.

#### 2.2.5. Sanger sequencing and ICE analysis

Sanger sequencing was performed using the BigDye™ Terminator v3.1 Cycle Sequencing Kit (Applied Biosystems), followed by capillary electrophoresis on an ABI 3500xl DNA Analyzer. Cycle sequencing reactions were set up in a 10 µL final volume containing 0.5–1.0 µL BigDye v3.1, 1.5 µL 5× Sequencing Buffer, 3.2 pmol sequencing primer, and 10–40 ng of purified PCR product, with nuclease-free water to volume. Thermal cycling was performed with an initial denaturation at 96°C for 1 min, followed by 25 cycles of 96°C for 10 s, 50°C for 5 s and 60°C for 4 min.

Prior to sequencing, PCR products were purified using ExoSAP-IT™ PCR Product Cleanup Reagent (Thermo Fisher Scientific to remove residual primers and dNTPs, according to the manufacturer’s instructions (37 °C for 15 min followed by enzyme inactivation at 80 °C for 15 min). Purified sequencing reactions were then subjected to capillary electrophoresis on the ABI 3500xl platform.

Resulting chromatograms were analysed using the ICE (Inference of CRISPR Edits) tool to estimate indel frequencies at the target locus and to infer the most likely insertion/deletion outcomes relative to the wild-type reference sequence, providing qualitative and semi- quantitative confirmation of genome editing outcomes.

### 2.3. Cas9 aggregation studies

#### 2.3.1. Samples for Cas9 aggregation studies

First, Cas9* solutions were prepared in buffered aqueous media to assess the presence of aggregation and to optimise the experimental techniques and analysis methods for its study.

Subsequently, series of independent Cas9* samples were prepared to study the potential effects of different parameters on Cas9’s aggregation. In all experimental series, the mean Cas9* concentration ranged from 0.5 – 2 nM, adjusting as necessary to maintain SMD conditions throughout the samples.

1. *Salt effect series*. In these series, we parted from the Cas9* stock solution (9 µg/µL in 50 % (v/v) in glycerol/water), diluting it using a solution of TRIS buffer and another concentrated solution of a salt (NaCl or MgCl_2_), also prepared in TRIS buffer. By diluting strongly the stock, solutions of Cas9* in TRIS buffer with different salt concentrations were obtained.
2. *Effect of sgRNA:Cas9 ratio*. In this series, the Cas9* stock solution was diluted using TRIS or PBS buffer to obtain salt-free or salt-containing samples, respectively. A separate sgRNA solution in the same buffer was prepared, and the two solutions were mixed to create samples with varying sgRNA:Cas9 ratios under salt-free or salt-containing conditions.
3. *Concentration series.* For this series, the Cas9* and Cas9 stocks were diluted using TRIS or PBS buffer, for the salt-free or the salt-containing samples, adding increasing amounts of Cas9 for the protein concentrated samples. In this way, up to 1.2 μM of Cas9 total concentration was obtained, while maintaining the Cas9* concentration constant at around 1 nM. To avoid any potential effects on solution viscosity and diffusion, a maximum glycerol concentration of 6.5 % (v/v) was reached in the highest Cas9 concentrated sample.

The aggregation studies for the *in vitro* cleavage experiments were carried out under the same conditions. Mixtures of Cas9* and the sgRNA A, also used in the cleavage assays, with a 1:1 molar ratio were prepared. The aggregation was measured in the three buffers of the cleavage experiments. The approximate Cas9* concentration was 0.5 nM for all samples.

#### 2.3.2. SMD/FCS measurements

The aggregation study was carried out using single-molecule fluorescence detection (SMD) and fluorescence correlation spectroscopy (FCS). The Cas9* concentration was chosen to be sufficiently low that, on average, only a single particle passes through the focal volume. Analysis of the detected photons allows the nature of the particle, monomer or aggregate, to be identified based on their different diffusive properties.

Single molecule fluorescence experiments were conducted using a custom-built setup. The light of a CW diode laser at 635 nm (Oxxius, LBX-638, FR) was coupled to a monomode optical fiber (Thorlabs, RB41A1, US), collimated (Schäfter&Kirchhof, 60FC-4-A7.5-01-DI, DE) and redirected by a dichroic mirror (Semrock, Brightline BS R635, US) and focused into the sample by a microscope objective (Olympus, UPLSAPO 60xW/1.20, water immersion) mounted in an inverted microscope (Olympus, IX71). The resulting fluorescence was collected by the same objective, directed through a pinhole (Thorlabs, Ø = 100 μm, US) and split into two beams by a nonpolarizing beamsplitter cube (Newport, 05BC17 MB.1, US). The fluorescence was separated from scattered light through a band-pass filter (Semrock, Brightline HC 679/41, US). Each of these beams was then focused onto an avalanche photodiode (MPD50CTC APD, Ø = 50 μm, MPD, IT). The signals from the detectors were processed and recorded by a time-correlated single photon counting (TCSPC) module (MultiHarp 150, PicoQuant, DE).

A 100 µL drop of each sample was deposited on clear flat bottom black polystyrene 96-well microplate. The measurements were collected at 25.0 ± 0.5 °C. An excitation laser power of *P* = 200 μW was selected, corresponding to a mean irradiance of *I*_0_/2 = 34.9 ± 0.4 kW cm^-2^. Even though a power series could not be performed to select the excitation power, photobleaching is unlikely due to low irradiances. Cy5 was used to calibrate the sample volume, being the same dye as the fluorescent label used in Cas9, using a diffusion coefficient of (3.6 ± 0.1) × 10^10^ m^2^ s^-1^ at 25 °C, based on the PicoQuant Application Note (19). A mean radial 1/e^2^ radius of W_*xy*_ = 0.604 ± 0.004 µm with a mean volume of *V* = 3.73 ± 0.09 µm^3^ was obtained for the focal volume for all experiments.

#### 2.3.3. SMD/FCS data analysis

A dedicated analysis framework, developed by our group, was applied to independently resolve and quantify monomeric and aggregated Cas9 species. In this approach, individual molecular transits are first extracted from the photon stream as fluorescence bursts, which are then characterized by their brightness, *I*_B_, and burst duration, *t*_B_, the latter reporting on the hydrodynamic size of the diffusing species. These burst-derived parameters are subsequently used to build the weights for the population separation, allowing us to isolate monomeric and aggregated Cas9 without requiring prior brightness knowledge, that we would need in standard FCS. From the resulting species-resolved populations, we determine diffusion properties and the fraction of aggregated protein using theoretical framework established by Fries *et al*. (20) and our previously developed aggregation models (10, 11, 21). Custom-made scripts developed in Python were used for this analysis and the *tttrlib* library was used to read the data from the TCSPC module (22). This method, that we called species-weighted Fluorescence Correlation Spectroscopy (swFCS), is briefly summarized in the Supplementary Information and will be described in detail in a forthcoming publication. Reported uncertainties for fitted parameters correspond to the estimated statistical standard deviations of the fits, with error propagation.

## 3. RESULTS

### 3.1. In vitro DNA cleavage efficiency

The *in vitro* cleavage assays were designed to analyse the effect of different parameters on DNA editing efficiency. In addition to the three selected protocols, which use different buffer types and compositions, the potential influence of the sgRNA type, the DNA substrate and the reaction time was also investigated.

Agarose gel (1 %) electrophoresis was performed for a qualitative determination of the genome editing output (Supplementary Figure S2), the intact DNA size and smaller fragments of sgRNA A and sgRNA C under different digestion conditions can be observed.

Three different sgRNAs (**Table 1** and **Table 2**) targeting cleavage sites A, B, and C (Supplementary Figure S1) were used at two reaction times (15 and 60 min) in two different DNA substrates (*ppifb* and *ppifb* WT). The *in vivo* activity of these sgRNAs was confirmed in CRISPR/Cas9-injected F0 embryos, which displayed mosaic editing patterns at the target sites as assessed by Sanger sequencing and ICE analysis (Supplementary Figure S3).

Capillary electrophoresis was used to quantify cleavage products based on fragment size and peak area (Supplementary Figure S4). In the *ppifb* control trace, two closely spaced full-length peaks were observed around ∼455 bp (apparent size), consistent with a heterozygous amplicon (Supplementary Figure S4A). In cleavage reactions, the full-length peak was accompanied by additional, guide-dependent shorter peaks together with a residual full-length peak (Supplementary Figure S4B–C). For quantitative analysis, peak areas were extracted using predefined size windows as explained in section 2.2.4. Based on the annotated cut positions, the expected cleavage fragments for the WT template are 191 bp (sgRNA A), 240 bp (sgRNA B) and 304 bp (sgRNA C), whereas the *ppifb* template is expected to yield fragments shifted by +4/5 bp (195/309 bp, respectively). Across traces, fragment sizes are reported as apparent sizes from capillary electrophoresis, which differed slightly from theoretical lengths (e.g., full-length peaks around ∼455 bp versus a 459-bp amplicon). The observed guide-dependent shorter peaks together with the residual full-length peak are consistent with partial cleavage of the substrate under these conditions.

Sanger sequencing revealed sequence variations within the amplified region of the *ppifb* template; however, most variants were located outside the PAM and protospacer sequences of the three sgRNAs and are therefore not expected to interfere with guide binding or cleavage. Specifically, for sgRNA A, a 4-bp insertion (CTGA) located 36 bp downstream of the predicted cut site and a single-nucleotide substitution (C>A) 43 bp downstream were detected. For sgRNA B, a single-nucleotide substitution located 2 bp upstream of the PAM was identified. For sgRNA C, a single-nucleotide substitution (C>G) and a single-nucleotide insertion (A) were detected within the protospacer sequence. These protospacer variants were present only in the *ppifb* template and therefore only in a subset of amplicon molecules, consistent with the mixed allelic composition of the heterozygous template.

**Figure 1** illustrates the influence of these parameters on cleavage efficiency. Cleavage efficiencies were comparable between *ppifb* WT and *ppifb* templates (**Figure 1A**), and only a modest trend towards increased cleavage was observed with longer incubation (60 min) (**Figure 1B**). However, significant variability was observed across sgRNAs under the conditions tested (**Figure 1C**). For subsequent analyses focused on buffer composition and Cas9 aggregation, sgRNA A was selected as a representative guide to minimise additional sources of variability.

**Figure 1.**
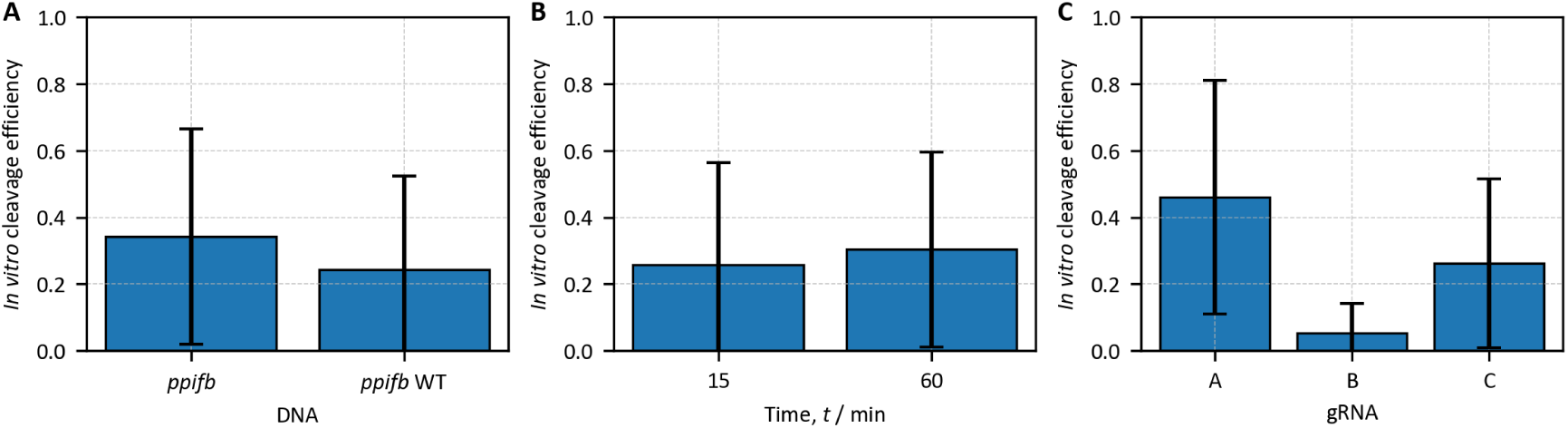
Effect of experimental factors on the *in vitro* editing efficiency. Bar plots show mean and standard deviation of editing efficiencies grouped by (A) DNA substrate, (B) incubation time, and (C) sgRNA type.

Therefore, to analyse the effects of buffer composition, subsequent analyses were restricted to reactions performed with sgRNA A, and data were pooled across DNA templates and reaction times because these factors did not have a major impact on cleavage efficiency under our conditions. The corresponding data, shown in **Figure 2**, demonstrate that cleavage efficiency is heavily affected by the buffer used in the reaction. This very significant effect could be due to the different composition of the buffers, potentially leading to different Cas9 aggregation levels. Therefore, we analyse the aggregation behaviour of Cas9 in these buffers below.

**Figure 2.**
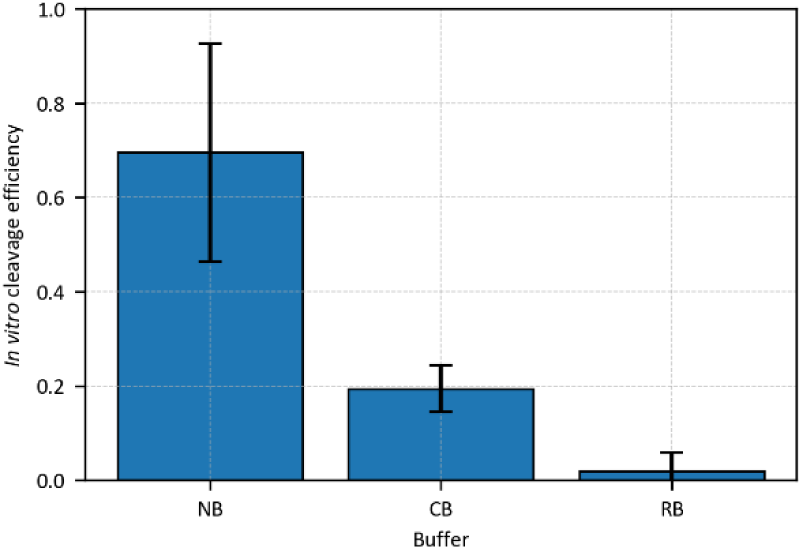
Effect of reaction buffers on the *in vitro* editing efficiency. Bar plots show mean and standard deviation of editing efficiencies.

### 3.2. Aggregation of Cas9 protein in aqueous solution

As a first experiment to assess the aggregation of Cas9 protein preparations, four independent Cas9* samples, each with a concentration ranging from 1 – 2 nM, were prepared in TRIS buffer (10 mM, pH 7.4), diluting strongly from the concentrated Cas9* stock (9 µg/µL in 50 % (v/v) water/glycerol).

Single molecule fluorescence measurements were carried out, revealing in all samples two distinct populations: one corresponding to monomers and the other to aggregates, when obtaining a 2D logarithmic heatmap of burst duration, *t*_B_, vs. count rate, *I*_B_, of the fluorescent bursts (see **Figure 3A**). We assign the populations in this way because of the burst duration: monomers diffuse faster through the focal volume, producing shorter fluorescent bursts, whereas aggregates are larger and diffuse more slowly, resulting in longer bursts. The populations are predominantly separated by the burst duration, *t*_B_, axis. Weights were calculated for each population, obtaining weighted marginal histograms for each axis. FCS was then applied to the fluorescence data, using a weighted correlation function based on the two populations, achieving a fluorescence correlation curve for each, as shown in **Figure 3B** (see the Supplementary Information for further explanation on the swFCS analysis). The correlation curves were fitted using equation SI.1, obtaining diffusional properties for each population, like the translational diffusion coefficient *D* and hydrodynamic radius *R*_H_ (equations SI.4 and SI.6). This method allows for a clear separation of both populations and enable their independent characterization within the same sample. The mean values for all four samples and for the monomer and aggregate populations are shown in **Table 3**. These results highlight the presence of heavily aggregated particles, since their particle size is around 50-fold higher than their respective monomers.

**Figure 3.**
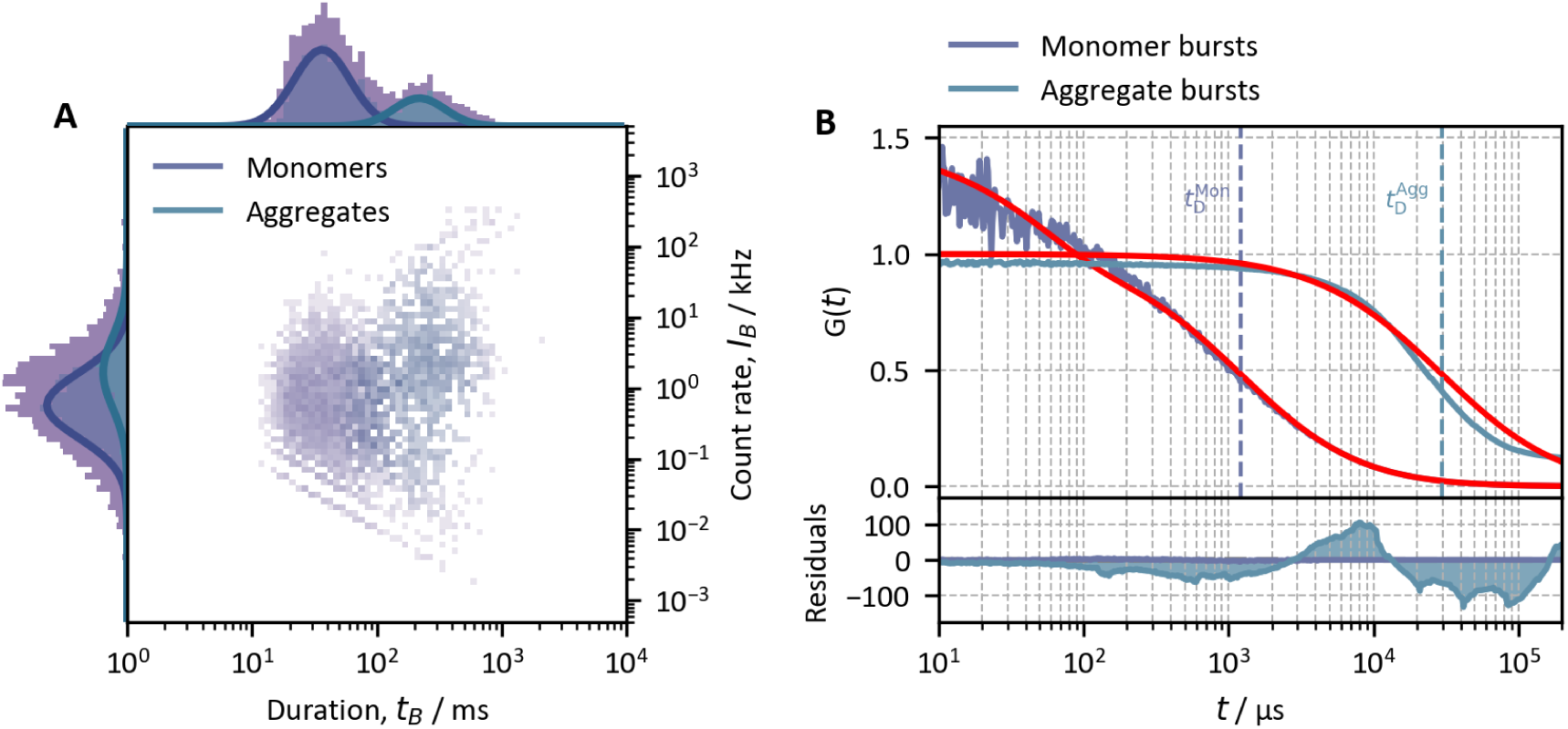
Left panel, A. 2D logarithmic heatmap of burst duration, *t*_B_, vs. count rate, *I*_B_, of the fluorescent bursts corresponding to one of the four samples. Weighted histograms for both monomer and aggregate populations are shown (see the Supplementary Information). A Gaussian function was fitted to obtain the mean duration and count rate for each population. Right panel, B. Normalized fluorescence correlation curves for the monomer and aggregate populations, obtained with a weighted correlation using their corresponding population weights (swFCS, see the Supplementary Information), along with the fit and weighted residuals of equation SI.1.

**Table 3.** Diffusional properties of monomeric and aggregated Cas9* in Tris buffer (10 mM, pH 7.4) at 25 °C.

| | $D / 10^{-12} \text{ m}^2 \text{ s}^{-1}$ | $R_H / \text{nm}$ |
| --- | --- | --- |
| Monomers | $66 \pm 2$ | $3.7 \pm 0.1$ |
| Aggregates | $1.37 \pm 0.03$ | $179 \pm 4$ |

With further analysis, quantifying the populations is also possible, through the amplitude of the FCS curves or directly through a photon count burst distribution analysis (see the Supplementary Information). Knowing the total concentration of Cas9 in the sample, the aggregation fraction, *γ*, can be calculated, which is the fraction of aggregated protein in the sample. All four samples showed a mean *γ* = 0.970 ± 0.001, highlighting the high aggregation level of Cas9 in aqueous solution (Tris buffer, 10 mM, pH 7.4). A mean aggregation number, *n̄*, meaning the mean number of monomers per aggregate, was calculated too, obtaining a mean number of *n̄* = 20 ± 5 monomers taking part in an aggregate.

### 3.3. Effect of salt, sgRNA and protein concentration on Cas9’s aggregation

Composition of the Cas9 samples was systematically modified to investigate the potential effects of different components. Salt concentration and cation charge effects were evaluated by adding different concentrations of NaCl or MgCl₂ to Cas9* samples. Then, the potential influence on aggregation of the ratio sgRNA:Cas9* and of the total Cas9 concentration were studied in both salt-free and salt-containing samples using TRIS (10 mM, no salt, pH 7.4) or PBS buffer (12 mM phosphate, 137 mM NaCl & 3 mM KCl, pH 7.4, *I* = 0.18 M), respectively.

Single-molecule fluorescence measurements were performed and analysed as described previously. Two populations, monomers and aggregates, were identified, and their respective weights calculated. The corresponding fluorescence correlation curves were generated and analysed to quantify the two species, obtaining the aggregation fraction for each sample. The mean aggregation number, *n̄*, was also calculated (see the Supplementary Information for further information and analysis explanations).

The main results from the whole analysis are shown in **Figure 4**, listed in **Table 4** and are described below:

1. *Salt effects*. Cas9* aggregates were observed to disaggregate with increasing concentrations of NaCl and MgCl₂, although the latter exhibited a much weaker effect. For the NaCl, not only the *γ* (aggregation fraction) but also the *n̄* (mean aggregation number) systematically decreases as salt concentration is increased. These findings indicate that salt promotes disaggregation by breaking down existing aggregates into smaller units, increasing the concentration of monomers in solution (see **Figure 4A** & **B**). In contrast, the addition of MgCl₂ resulted in a less pronounced decrease in aggregation, with the mean aggregation number remaining very high and relatively constant (**Figure 4B** & **Table 4**).
2. *sgRNA effect*. The increasing ratio of sgRNA:Cas9 in the Cas9* aggregated samples is observed to increase the aggregation fraction, *γ*, in the titration series in PBS and the size of the aggregates *n̄*, in both buffers. These observations imply that sgRNA increases the stability of the aggregated particles of Cas9*, inducing aggregation. The effects in the Tris titration as not as apparent in the aggregation fraction, due to the starting point being much more aggregated. All results are shown in **Figure 4C** & **D**. It must be noted that the starting point of aggregation of the samples in PBS buffer matches the aggregation level observed for the corresponding NaCl concentration (around 137 mM) in **Figure 4A**.
3. *Concentration effect*. Increasing the protein concentration results in higher aggregation fractions both in PBS and in Tris buffers. However, smaller changes were observed in Tris buffer, where aggregation fractions were already high at low Cas9 concentrations. The aggregation fraction of these concentration series is shown in **Figure 4E**.

**Figure 4.**
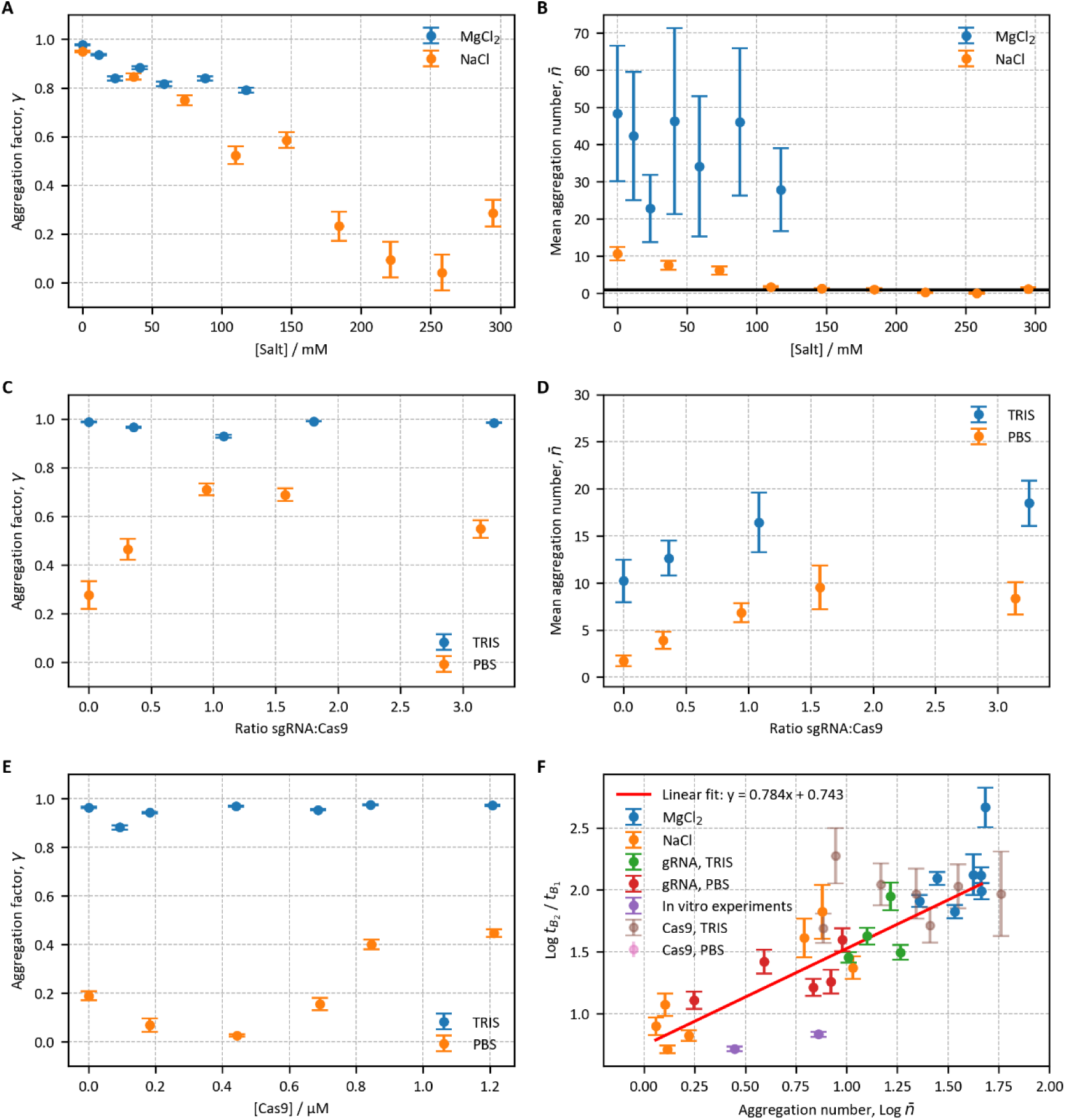
Left panels: A, C & E. Aggregation fraction, *γ*, of Cas9* samples with increasing amounts of salt (A), sgRNA:Cas9 molar ratio (C) and Cas9 concentration (E). The average Cas9* concentration was approximately 1 nM in all samples. Different salts, like MgCl_2_ and NaCl, or different conditions, like TRIS and PBS buffer, are shown in different colours. Right panels: B & D. Mean aggregation number, *n̄*, as a function of salt concentration (B, the horizontal black line represents *n̄* = 1) or sgRNA:Cas9 molar ratio (D). Right panel: F. Double logarithmic plot of the ratios of the aggregate and monomer burst duration (*t*_B_2__ /*t*_B_1__) of Cas9* versus the mean aggregation number, *n̄*. A linear fit is presented, whose slope showcases the shape factor, *ν*, which indicates the shape of the diffusing particle (equation SI.15). The different experimental series are also presented in different colours. The Cas9 concentration series are also presented even though they were not introduced in the fit.

**Table 4.**
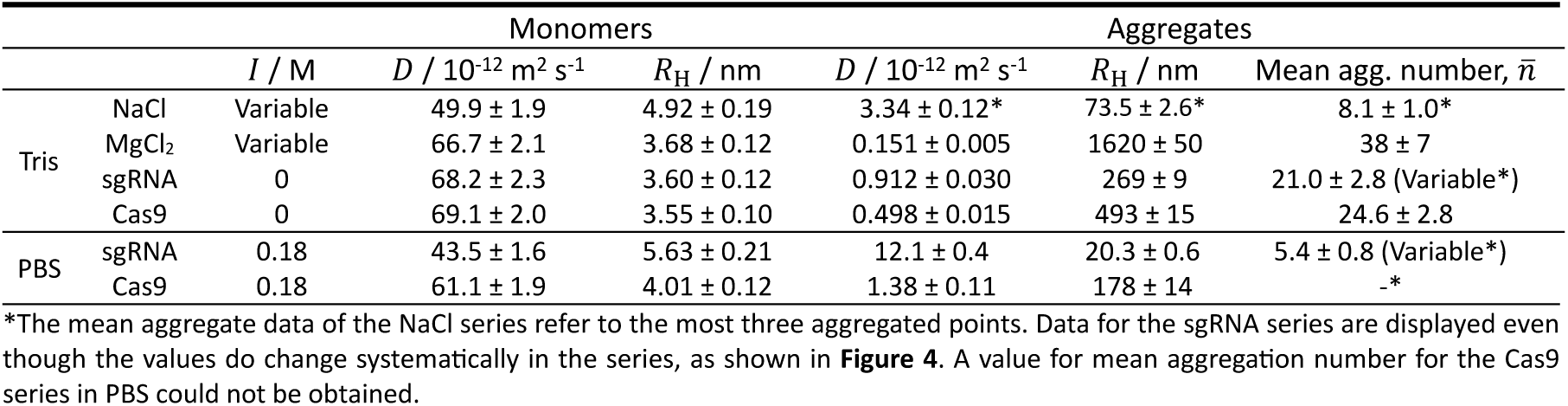
Mean diffusional properties of monomeric and aggregated Cas9* for the different series at 25 °C.

A shape index, *νν*, which provides information about the shape of the diffusing particle, (*νν* = 1/3 for homogeneous spheres, *νν* = 1/2 for random coils, *νν* = 1 for rigid rods), could also be calculated for the aggregates of the salt and sgRNA series, with the *in vitro* cleavage aggregation experiments (detailed analysis given in the Supplementary Information). A shape index of *ν* = 0.8 ± 0.1 was obtained from the linear relation between the ratio of the aggregate and monomer burst durations (*t*_B_2__ /*t*_B_1__) and the mean aggregation number, *n̄*, shown in **Figure 4F**.

### 3.4. Aggregation of Cas9 under in vitro cleavage conditions

The aggregation behaviour of Cas9 in the three buffers used in *in vitro* experiments was studied. Cas9 concentration was fixed at 0.5 nM across all samples. sgRNA A was included to simulate the *in vitro* experiment, with 1:1 molar ratio, just like the cleavage assays experiments. The same analytical procedure was applied in all cases. The aggregation fraction, *γ*, of these samples is shown in **Table 5**. We can observe that Cas9 does not aggregate in the NEBuffer (NB). In contrast, in the Reaction Buffer (RB) and in the Cleavage Buffer (CB) Cas9 exhibits high aggregation levels, with aggregate sizes comparable to those observed under other experimental conditions (**Table 3** and **Figure 4F**).

**Table 5.** Aggregation fraction, *γ*, and mean aggregation numbers, *n̄*, of Cas9* samples in the different reaction buffers of the *in vitro* editing assays.

|  | NEBuffer (NB) | Cleavage Buffer (CB) | Reaction Buffer (RB) |
| --- | --- | --- | --- |
| Aggregation fraction, $\gamma$ | 0 | $0.53 \pm 0.03$ | $0.88 \pm 0.01$ |
| Mean aggregation number, $\bar{n}$ | – | $2.8 \pm 0.8$ | $7 \pm 2$ |

## 4. DISCUSSION

### 4.1. Influence of experimental parameters on in vitro cleavage efficiency

Before addressing the role of Cas9 aggregation, it is important to briefly discuss the main trends observed in the *in vitro* cleavage efficiency assays (**Figure 1** & **Figure 2**). Under the conditions tested, no significant effect of the DNA substrate was detected, as comparable cleavage efficiencies were obtained for the *ppifb* WT and *ppifb* templates. Similarly, extending the reaction time from 15 to 60 min produced only a modest increase in cleavage efficiency, indicating that reaction time is not a limiting factor within this range. These results suggest that Cas9 cleavage is largely independent of substrate heterogeneity and incubation time under our experimental conditions.

In contrast, the choice of sgRNA had a measurable effect on cleavage efficiency. Among the three guides tested, sgRNA B consistently exhibited lower cleavage efficiency than sgRNAs A and C. Although some variability was observed between sgRNAs A and C, the differences between these two guides were not statistically significant with the data available. However, only the data obtained with sgRNA A were selected to compare the effect of the reaction buffers to minimise additional sources of variability.

Cleavage efficiencies varied dramatically depending on buffer composition, with the NB yielding the highest activity and the RB the lowest, a negligible activity. This strong buffer dependence cannot be explained by differences in DNA substrate, reaction time or sgRNA alone, and instead points to a dominant influence of the physicochemical environment on Cas9 functionality. As discussed below, these differences closely parallel the distinct aggregation states of Cas9 observed in each buffer, suggesting that buffer-dependent modulation of Cas9 aggregation is a key factor underlying the observed variations in cleavage efficiency.

### 4.2. Cas9’s aggregation behaviour

Our SMD/FCS experiments of Cas9* in buffered aqueous solution show clearly the presence of aggregates in equilibrium with monomers (**Figure 3**), with a very high aggregation fraction close to unity. The value for the hydrodynamic radius of the monomeric Cas9 (**Table 3**) is of the order of those reported in the literature for the same species, but significantly lower. Manzano *et al*. (6) recorded a value of 5.15 ± 0.75 nm for the monomeric Cas9 using Dynamic Light Scattering (DLS), but in slightly different conditions (20 mM Tris, 300 mM NaCl & pH 8.0). Similar results were obtained by Nguyen and coworkers (7), that recorded a value around 5 – 7.5 nm for the Cas9 monomer, also in similar conditions (10 mM Tris, 150 mM KCl & pH 7.4).

Our observed hydrodynamic radius for the monomeric Cas9 is consistent with predictions based on the polypeptide chain length. Using the empirical relationship *R*_H_ = (4.75 ± 1.11) *N*^(0.29±0.02)^ Å (23), where *N* is the number of amino acids (*N* = 1368 amino acids (24)), the expected value for the monomeric Cas9 is 3.9 ± 0.1 nm, which aligns perfectly with our experimentally observed value 3.7 ± 0.1 (**Table 3**), a value that was obtained by measuring aqueous solutions of Cas9 (where aggregates appear). However, the monomer diffusion coefficient and hydrodynamic radius can also be obtained from some of the titration experiments, where higher fractions of monomeric Cas9 was available. That is the case of the NaCl series, where the aggregation fraction remained low in the presence of high salt concentrations. In this way, the mean value for that titration series was 4.9 ± 0.2 nm (**Table 4**), which aligns better with Manzano (6) and Nguyen (7) results. This discrepancy could be attributed to the much more precise value for the monomer obtained in the latter case, because of the higher fraction of monomers in the samples, or to a potential conformational change of the protein induced by salts, since the literature values were measure in the presence of salts too. This could be the case since the size of the monomer in the presence of sgRNA is different for the case of Tris and PBS buffer, as appear in **Table 4**. In this case, we would be observing the size of the complex sgRNA:Cas9. This possible change in conformation is not observed in the case of the MgCl_2_ titration, but this could be due to the fact that Mg^2+^ is an enzyme cofactor and binds to the protein (25, 26).

Regarding aggregated Cas9, Jain and coworkers (8) also studied its aggregation behaviour. They recorded an increase in Cas9 size with rising pH (ranging from 5 to 8) and observed a value around 400 nm for the Cas9 at pH 7, also measured with DLS. Their observed value is significantly higher than our mean value of the aggregates, however, their Cas9 concentration was much higher (200 nM). They hypothesised that, as Cas9 approaches its isoelectric point (theoretical *pI* = 9) and loses its positive charge, its ability to bind to negatively charged compounds like sgRNA decreases. At the same time, the loss of net charge reduces electrostatic stabilization by ions in the aqueous medium and weakens inter-monomer electrostatic repulsion, thereby decreasing colloidal stability and promoting aggregation. What’s more, Nguyen and coworkers (7) also reported that Cas9 tended to precipitate, with solubility increasing upon the addition of sgRNA, but observed the formation of aggregates when sgRNA was added to Cas9 through DLS, with a mean size of 100 nm for these aggregates. Our measurements showed high variability of the size of Cas9 aggregates, as displayed in **Table 3** & **Table 4**, with values that also include the ones found in the literature.

In our analysis we obtain also an interesting information regarding the shape of Cas9 aggregates. As shown in **Figure 4F**, a shape index of *ν* = 0.8 for the aggregates is estimated from the linear relation between the ratio of the aggregate and monomer burst durations (*t*_B_2__ /*t*_B_1__) and the mean aggregation number, *n̄*. The value obtained shows that the aggregates have an elongated, cylindrical shape. Moreover, the shape is independent of the aggregates’ size, since the data from all series, with different aggregation numbers (**Table 4**), fit satisfactorily the same straight line.

As for the salt, sgRNA:Cas9 ratio and protein concentration aggregation series, some results can be inferred from the data:

1. *Salt effects*. Increasing the concentration of salt appears to promote the disaggregation of large Cas9 protein particles. This likely occurs through the breakdown of larger aggregates into their monomeric subunits, as seen with NaCl (**Figure 4B)**. The positively charged Cas9 monomers become more stable in solutions with higher ionic strength, shifting the aggregation equilibrium toward the monomeric form. In contrast, MgCl_2_ was less effective in promoting disaggregation compared to NaCl. This may be due to the presence of larger aggregates in the initial MgCl_2_ sample, which could be more resistant to disaggregation or exhibit irreversible behaviours. What’s more, Mg^2+^ is a cofactor for the Cas9 enzyme (25, 26), which means that the metal binds to the protein and is necessary for enzymatic activity. The binding of the metal to the protein could help to increase its positive charge, making the aggregates more stable and harder to break down by the salt.
2. *Effect of sgRNA:Cas9 ratio*. Although it may seem counterintuitive that sgRNA, a negatively charged molecule with a high affinity for the Cas9 monomer, could promote aggregation, it is possible that sgRNA interacts with positively charged outer regions of larger Cas9 aggregates. This interaction could partially neutralize their surface charge, reduce colloidal stability and promote aggregate growth, as we can observe in **Figure 4D**, where the aggregation number increases for both titrations. According **Figure 4C**, in the absence of salt Cas9 is almost completely aggregated, and no significant effect of the sgRNA:Cas9 ratio is observed. In contrast, in PBS a clear dependence on the sgRNA:Cas9 ratio emerges: the aggregate fraction decreases at low sgRNA:Cas9 ratios, while remaining high for ratios larger than 1:1. These data suggest that aggregation becomes stabilized once a 1:1 sgRNA:Cas9 ratio is reached, beyond which further addition of sgRNA does not significantly reduce aggregation, although a slight trend toward monomerization is observed at higher ratios. This behaviour is consistent with the results reported by Jain and coworkers (8), who observed enhanced Cas9 aggregation when approaching its *pI*, as well as with the observations of Nguyen *et al*. (7), who reported aggregation upon sgRNA addition but also partial dissolution of Cas9 at high sgRNA:Cas9 ratios.
3. *Concentration effect*. The aggregation fraction, *γ*, increased with protein concentration in both PBS and Tris buffers, making the changes more apparent in the case of the PBS, since *γ* starts at lower values. A similar effect of the sgRNA could happen when adding unlabelled Cas9. For the case of the samples in Tris buffer, the added Cas9 seemed not to interact with the preformed aggregates, since no changes in the fluorescence intensity of the aggregates is observed. This suggests that the pre-formed aggregates are highly stable and possess a strong positive charge, preventing the unlabelled monomers from incorporating into the aggregates. Instead, the added Cas9 could likely form its own unlabelled aggregates, either with other unlabelled monomers or with monomeric Cas9*. However, these new aggregates would exhibit a much lower degree of labelling, which would be eclipsed by the brighter pre- formed aggregates and not seen in our measurements. For the case of the concentration series with PBS buffer, the aggregation fraction, *γ*, did increase.

### 4.3. Efficiency vs. aggregation

Based on the results of the aggregation studies, we can state that the different aggregation fraction obtained in the *in vitro* cleavage conditions can be explained by the buffer compositions of the solution used (**Table 5**). Among the tested conditions, the NB buffer exhibited the lowest level of aggregation. This observation is consistent with the fact that NB and PBS share the same ionic strength (*I* = 0.18 M, see sections 2.1 and 2.2.3 for buffer composition), but NB also contains BSA. At pH 7.9, BSA is negatively charged and can interact with Cas9, stabilizing its positively charged monomers and thereby promoting disaggregation. In contrast, the CB and RB buffers have lower ionic strengths (*I* = 0.14 M and *I* = 0.12 M, respectively), which account for the higher levels of aggregation observed under those conditions.

Now we analyse the potential relationship between Cas9 aggregation and the *in vitro* cleavage efficiency. To facilitate the discussion of this behaviour we have summarized in **Table 6** the composition of the buffers and their ionic strength (see section 2.2.3 for more details), the aggregation fraction and mean aggregation number, and the cleavage efficiencies for each buffer. As observed, the values of aggregation fraction and aggregation number diminish with increasing cleavage efficiencies. As explained before, the significant variations in efficiency can only be due to the reaction buffers. So, we can infer that there is an inverse correlation between these variables. To investigate further, we plotted the editing efficiency as a function of the aggregation fraction, *γ*, in **Figure 5**. A linear regression was then performed to assess the inverse correlation between Cas9 aggregation and its cleavage efficiency. A high inverse correlation was obtained (the slope being −0.8 ± 0.1).

**Figure 5.**
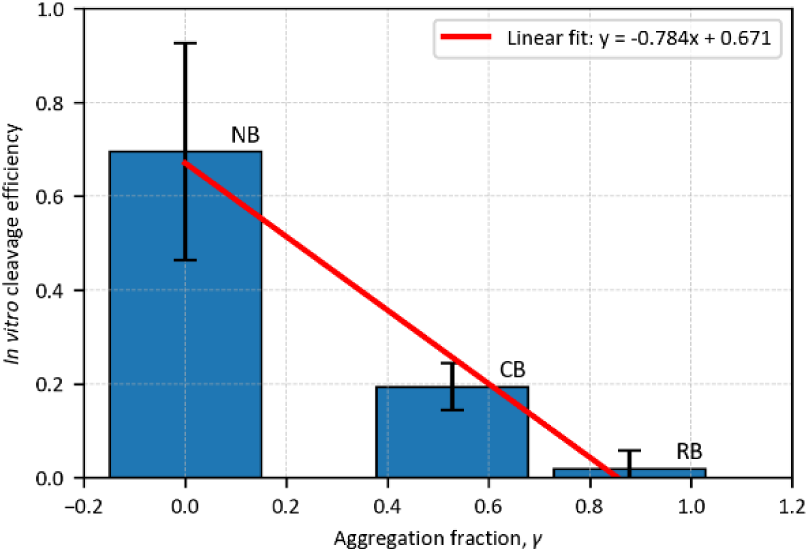
Effect of reaction buffers on the *in vitro* editing efficiency. The efficiency data corresponding to the different reaction buffers (also presented in **Figure 2**) is plotted against the corresponding aggregation fraction, *γ*. A linear fit is performed to demonstrate the inverse correlation between Cas9’s aggregation and its *in vitro* cleavage efficiency.

**Table 6.** Aggregation fraction, *γ*, mean aggregation number, *n̄*, and cleavage efficiency of Cas9* for the buffers used for the *in vitro* assays, with its composition and ionic strength, *I*.

| Buffer | Composition | $I / M$ | Agg. fraction, $\gamma$ | Mean agg. number, $\bar{n}$ | Cleavage efficiency / % |
| --- | --- | --- | --- | --- | --- |
| NEBuffer (NB) | NaCl, MgCl<sub>2</sub>,<br>Tris-HCl, BSA;<br>pH 7.9 | 0.18 | 0 | – | $0.7 \pm 0.3$ |
| Cleavage Buffer (CB) | KCl, MgCl<sub>2</sub>,<br>Tris-HCl,<br>glycerol, DTT;<br>pH 7.5 | 0.12 | $0.53 \pm 0.03$ | $2.8 \pm 0.8$ | $0.19 \pm 0.05$ |
| Reaction Buffer (RB) | NaCl, MgCl<sub>2</sub>,<br>HEPES, EDTA | 0.14 | $0.88 \pm 0.01$ | $7 \pm 2$ | $0.02 \pm 0.03$ |

Aggregation impairs the protein functionality while also reducing the amount of functional monomeric protein in solution. This outcome highlights that that careful control of protein solubility is crucial for maintaining both activity and effective downstream applications. We suggest that experimental conditions should be optimised to keep Cas9 fully dissolved, not only to improve its functionality but also in delivery assays where the protein is encapsulated, where aggregation could impede protein loading and internalisation. Similarly, *in vivo* applications may be affected, as aggregated protein could trigger undesired immune responses (5).

## 5. CONCLUSIONS

Our study shows a robust inverse correlation: greater Cas9 aggregation consistently leads to lower genome-editing *in vitro* efficiency, regardless of sgRNA type, target DNA, or incubation time. By combining single-molecule fluorescence approaches with quantitative cleavage assays, we show that aggregation significantly reduces Cas9 activity and that higher aggregation levels consistently yield lower cleavage efficiency.

Our results identify buffer composition, ionic strength, protein concentration and sgRNA presence as key factors modulating Cas9 aggregation. Conditions that increase ionic strength or include stabilizing components effectively reduce aggregation and preserve a higher population of functional, monomeric Cas9, leading to improved cleavage performance.

Together, these findings establish Cas9 aggregation as a major physicochemical factor limiting *in vitro* genome-editing efficiency. Understanding and controlling aggregation is therefore essential for the rational optimization of CRISPR/Cas9 experimental protocols and for the development of more efficient and reliable genome-editing and delivery strategies.

## 6. ACKNOWLEDGEMENTS

A.S. thanks the Xunta de Galicia for her predoctoral contract. D.C. thanks the Xunta de Galicia for his predoctoral contract “Campus de Especialización Campus Terra”.

## 7. AUTHOS CONTRIBUTIONS

- **Diego Cora**: Investigation, Data Curation, Formal analysis, Software, Visualization, Methodology, Writing – original draft.
- **Ana Seijas**: Investigation, Data Curation, Formal analysis, Writing – original draft.
- **Wajih Al-Soufi**: Conceptualization, Methodology, Project administration, Resources, Supervision
- **Laura Sánchez**: Conceptualization, Project administration, Supervision
- **Álvaro J. Arana**: Conceptualization, Investigation, Data Curation, Formal analysis, Methodology, Writing – original draft, Funding Acquisition, Project Administration, Resources, Supervision
- **Mercedes Novo**: Conceptualization, Writing – Review & Editing, Methodology, Funding Acquisition, Project Administration, Resources, Supervision

