## Supplementary Information for "Critical role of Cas9 aggregation on *in vitro* DNA cleavage"

### Index

### 1. SUPPLEMENTARY DATA

**Supplementary Table S1**. Primers used for amplification and fragment analysis of the *ppifb* amplicon.

| | Sequence (5'→3') | Length / nt | $T_m$ / °C | GC % | Product length / bp |
|---|---|---|---|---|---|
| *ppifb*-F | 6-FAM-ATCGCCCATATCGCAATGCT | 20 | 60.32 | 50.00 | 459 |
| *ppifb*-R | CCCTTCGCTGATTCGTTAGC | 20 | 59.07 | 55.00 | |

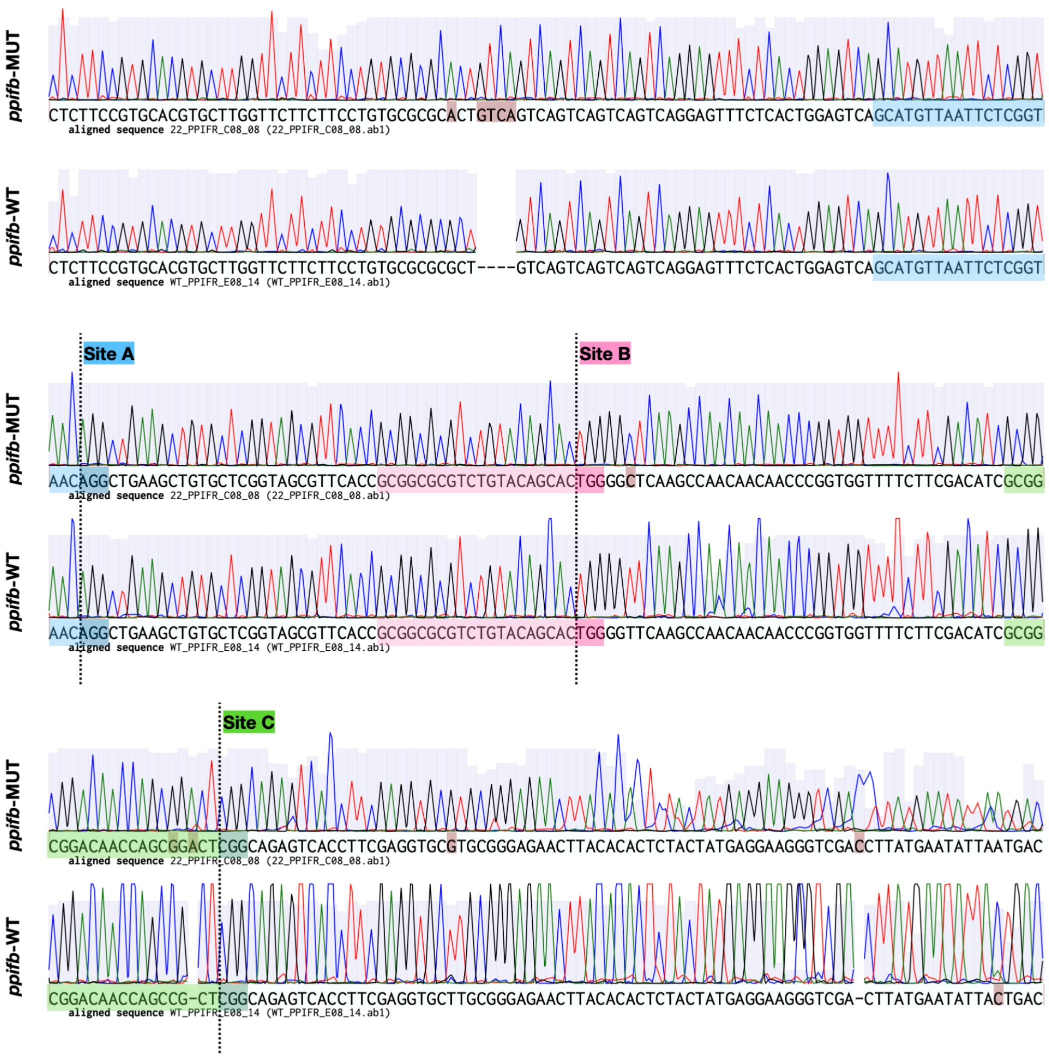

**Supplementary Figure S1**. Sanger sequencing chromatograms of the *ppifb* WT and *ppifb* (shown as *ppifb*-MUT in the figure) amplicons used as substrates in the *in vitro* cleavage assays, and mapping of the three CRISPR/Cas9 target sites. Representative forward and reverse reads were aligned to the reference *ppifb* amplicon, and the target sites for sgRNA A, sgRNA B and sgRNA C are indicated as Site A, Site B and Site C (vertical dashed lines; PAM highlighted). For the *ppifb* WT template, no sequence differences were detected within the protospacer or primer-binding regions across the three target sites. For the *ppifb* template, small sequence differences within the amplified region were observed between alleles; these were located outside the guide-binding regions for sgRNA A and sgRNA B, whereas sgRNA C showed an allelic variant within the protospacer sequence. Chromatograms were visualized and annotated using Benchling.

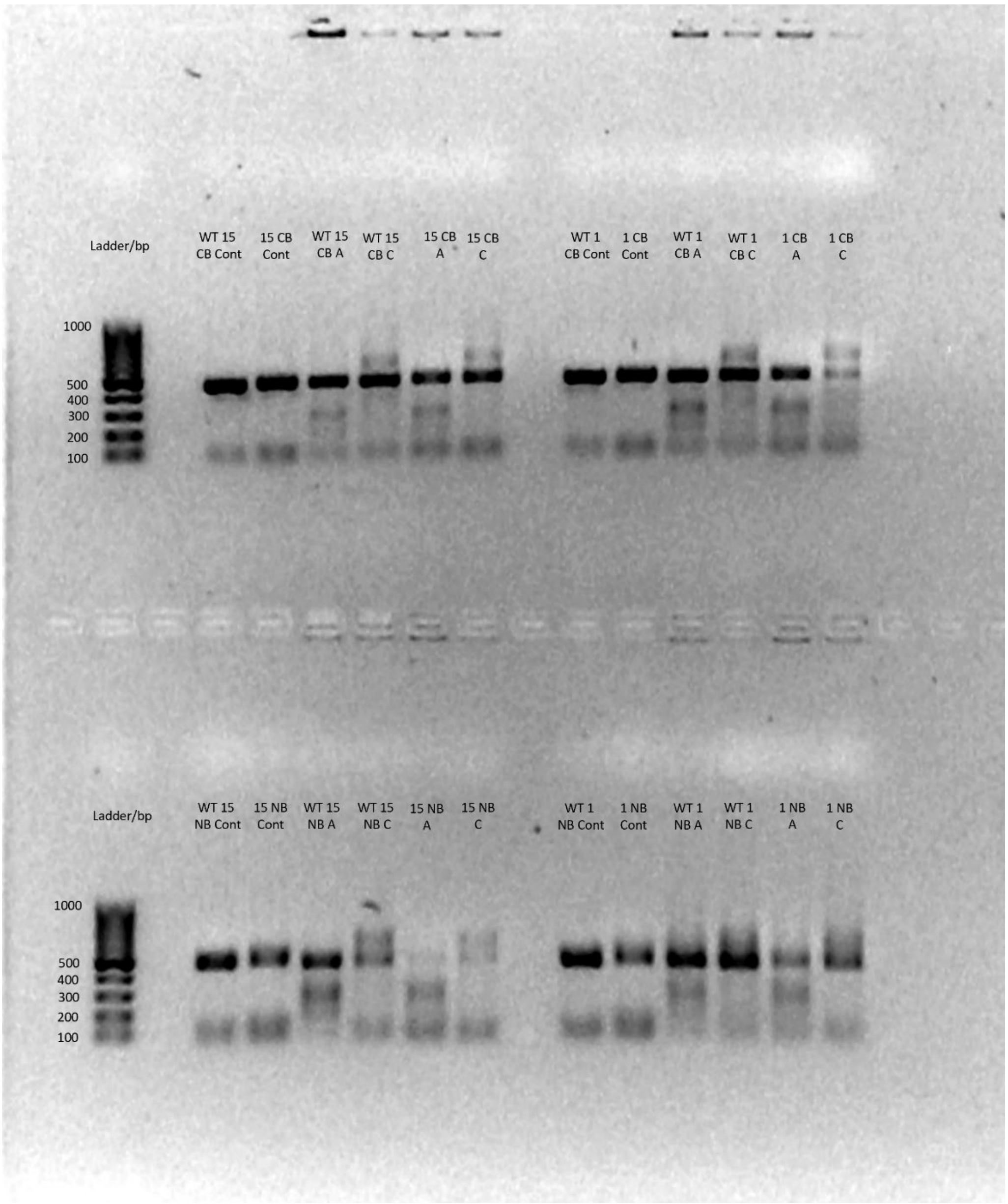

**Supplementary Figure S2.** Agarose gel (1%) electrophoresis was performed to analyse cleavage assays. Different samples of *ppifb* digested gene were loaded onto the gel. GeneRuler 100 bp DNA Ladder was loaded in the first column as a reference. The gel shows different samples of *ppifb* and *ppifb* WT, under two digestion times (15 minutes or 1 hour), using two different buffers (CB in the upper row or NB in the lower row) and two different gRNA (A or C). Note that control samples, were the digestion was performed neither with sgRNA or Cas9, were also loaded in the gel. DNA size is evident specially in control samples with bands around 500 bp. Smaller bands are clearly visible in cleavage assays performed with buffer NB, especially with sgRNA A where the band of non-cut DNA shows lower intensity. Some smaller bands are also observed in cleavage assays performed with buffer CB although with lower intensity.

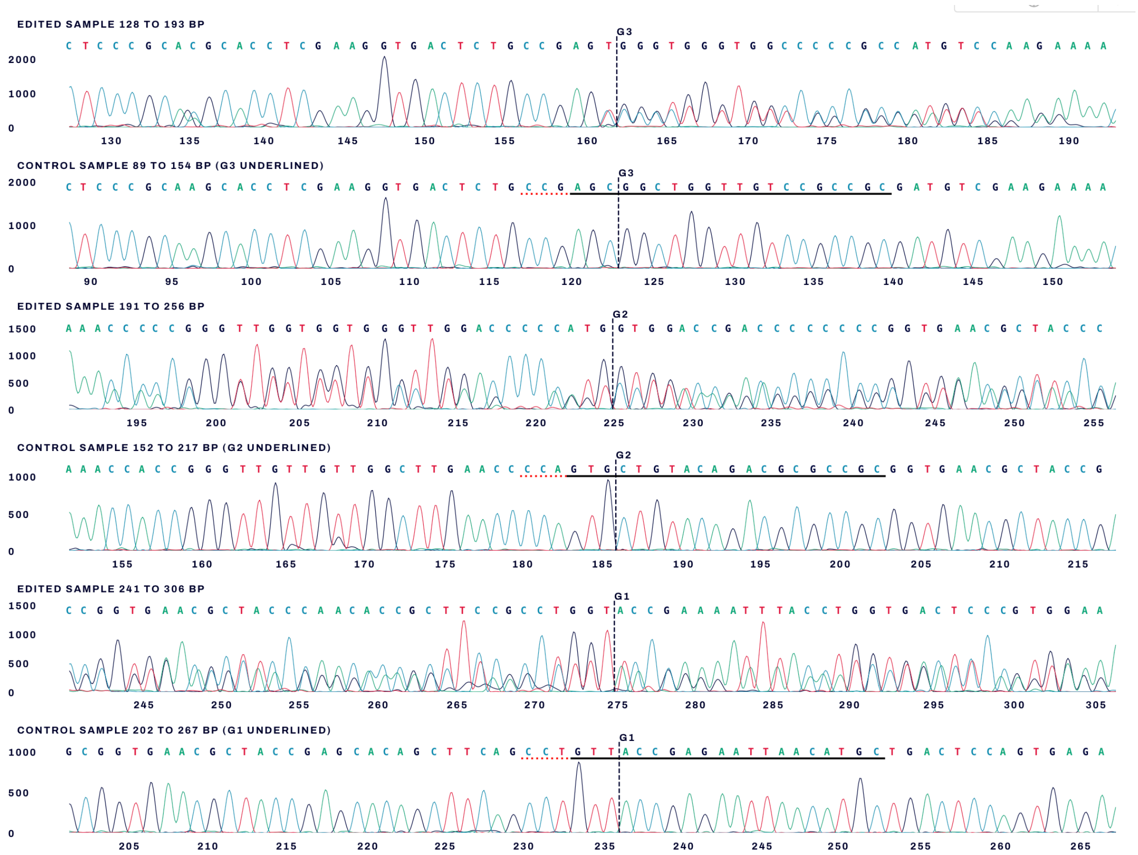

**Supplementary Figure S3.** Sanger sequencing traces from ICE analysis comparing edited samples (F0 individual, top panels) and the corresponding wild-type control (bottom panels) for the *ppifb* amplicon. The positions of the three CRISPR/Cas9 target sites are indicated by dashed vertical lines and labelled G1, G2 and G3, corresponding to sgRNA A, sgRNA B and sgRNA C, respectively. For each region, the chromatograms show sequence differences between the F0 and WT samples that are compatible with mosaic CRISPR editing, confirming that the three guides are able to induce DNA variants at the expected cut sites.

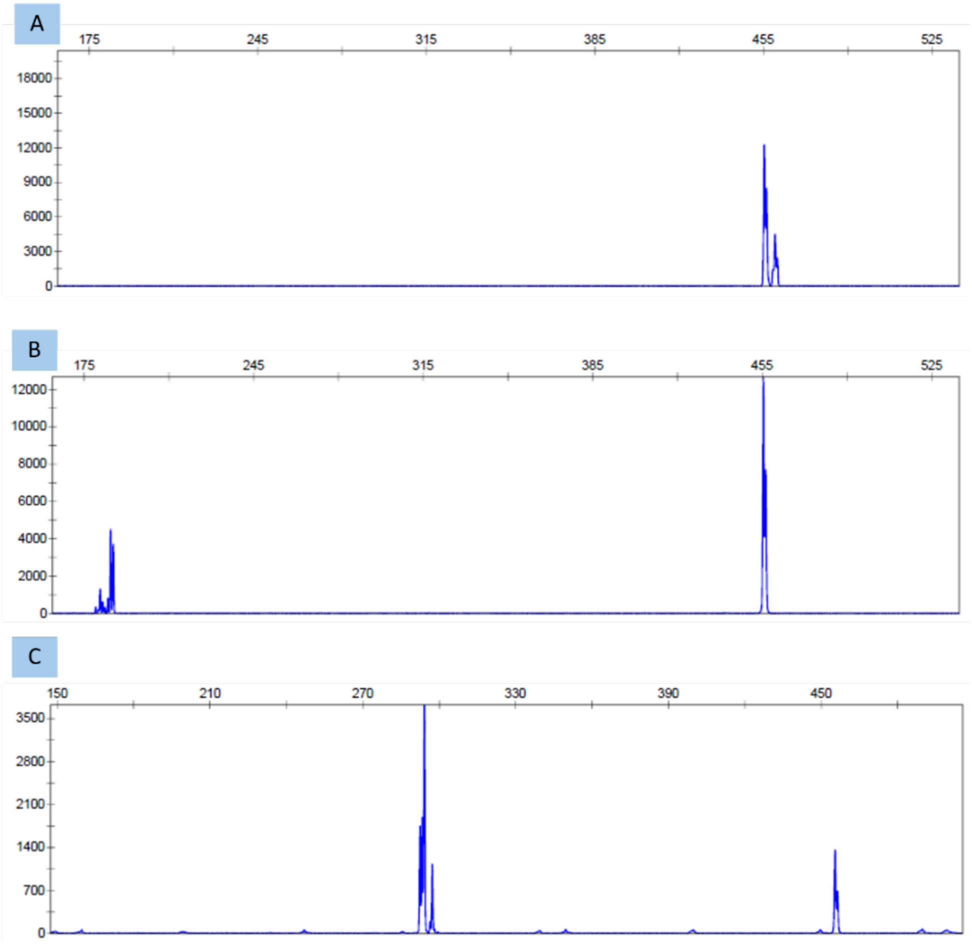

**Supplementary Figure S4**. Chromatograms obtained with ABI 3500xL Genetic Analyzer (Applied Biosystems) and GeneMapper™ Software v4.1 after cleavage assays. **A** *ppifb* control chromatogram whit CB buffer, two peaks around 455bp are visible which represent the heterozygosis of the mutant DNA. **B** *ppifb* WT with NB buffer and with one 60 min of cleavage assay time and RNA A. Several peaks are easily observed at 180bp as a result of the cleavage assay with RNA A but also a peak at 455bp of non-cuted DNA is visible. **C** *ppifb* WT with NB buffer and with 60 min of cleavage assay time and RNA C. Several peaks are easily observed at 290bp as a result of the cleavage assay with RNA C but also a peak at 455bp of non-cut DNA is visible.

### 2. SMD/FCS DATA ANALYSIS OVERVIEW

Single-molecule fluorescence data were analysed using a novel approach developed by our group, called swFCS, to independently resolve and quantify the two populations observed in the experiments, monomers and aggregates (see Figure 3 in the main article). This method, which will be described in detail in a forthcoming publication, builds upon the theoretical framework established by Fries *et al.* (1) for single-molecule quantification and extends our previously developed aggregation models (2–4).

#### *2.1. Burst detection and preprocessing*

Photon arrival times ($t_i$) were converted into interphoton times ($\Delta t_i = t_{i+1} - t_i$). A Lee filter ($m$ = 10, $\sigma_0 = 10$) was applied to smooth random fluctuations (5). Fluorescent bursts were identified using dual thresholds for background and fluorescence intensity. Each burst was characterized by:

- **Duration** ($t_{\mathrm{B}}$): time between first and last photon,
- **Photon count** ($C_{\mathrm{B}}$), and
- **Intensity** ($I_{\mathrm{B}} = C_{\mathrm{B}}/t_{\mathrm{B}}$).

These parameters were used to infer molecular brightness and diffusion behaviour.

#### *2.2. Weight-based population separation*

Monomeric and aggregated species were distinguished using statistical weighting based on burst duration and intensity. Logarithmic histograms of $t_B$ and $I_B$ were fitted with Gaussian functions to generate one-dimensional weights, which were then combined into two-dimensional weights, for each population. Weighted histograms allowed to obtain population-specific parameters, like mean intensity and duration (see Figure 3 in the main article).

#### *2.3. Species-weighted Fluorescence Correlation Spectroscopy (swFCS)*

To isolate the contribution of each population, photon events were weighted according to the burst-based weights prior to autocorrelation analysis, obtaining separate correlation curves for each population, which we called species-weighted Fluorescence Correlation Spectroscopy (swFCS). To improve correlation quality, burst regions were symmetrically expanded 1 s to capture fluctuations exceeding the diffusion timescale (6, 7).

Weighted mean count rates were computed as the inverse of the weighted mean interphoton time. Background signal corrections were corrected using the equation $\chi^2 = (1 + \langle b \rangle / \langle f \rangle)^{-2}$, (1, 8). In this way, $N$ can be corrected multiplying by the inverse of $\chi^2$ and $I$ by subtracting the background count rate $\langle b \rangle$. From this point forward, $N$ and $I$ will be considered as corrected values in the subsequent analysis.

The fluorescence correlation curves were analysed by fitting a model that includes two diffusion components and two triplet-state or reaction contributions:

$$G(t_{\mathrm{C}}) = \frac{1}{N}\Big(R \cdot G_{\mathrm{D}_1}(t_{\mathrm{C}}) + (R-1) \cdot G_{\mathrm{D}_2}(t_{\mathrm{C}})\Big) \cdot G_{\mathrm{T}_1}(t_{\mathrm{C}}) \cdot G_{\mathrm{T}_2}(t_{\mathrm{C}}) + b_0 \quad (1)$$

where:

- $N$ is the apparent number of molecules in the observation volume,
- $R$ is the contribution of the first diffusion correlation time to the correlation curve,
- $b_0$ the baseline value for the correlation curve,
- $G_{\mathrm{D}_i}(t_{\mathrm{C}})$ describes the diffusion term for species $i$ = 1, 2,
- $G_{\mathrm{T}_i}(t_{\mathrm{C}})$ models the triplet-state relaxation or other photophysical processes.

The diffusion and triplet terms are defined as:

$$G_{\mathrm{D}_i}(t_{\mathrm{C}}) = \left(1+\frac{t_{\mathrm{C}}}{t_{\mathrm{D}_i}}\right)^{-1}\left(1+\frac{t_{\mathrm{C}}}{w^2\, t_{\mathrm{D}_i}}\right)^{-1/2} \tag{2}$$

$$G_{\mathrm{T}_i}(t_{\mathrm{C}}) = A_{\mathrm{T}_i}\exp\left(-\frac{t_{\mathrm{C}}}{t_{\mathrm{T}_i}}\right) \tag{3}$$

where:

- $t_{\mathrm{D}_i}$ is the diffusion correlation time of species $i$ diffusing through the focal volume,
- $w$ is the aspect ratio of the focal volume,
- $A_{\mathrm{T}_i}$ is the amplitude of the term,
- $t_{\mathrm{T}_i}$ is the corresponding relaxation time.

The diffusion time, $t_{\mathrm{D}_i}$relates to the translational diffusion coefficient, $D_i$, through:

$$t_{\mathrm{D}_i} = \frac{w_{\mathrm{xy}}^2}{4D_i} \tag{4}$$

where $w_{\mathrm{xy}}$ is the axial radius of the focal volume. Measuring a substance with a known $D$, the $w_{\mathrm{xy}}$ can be calibrated and the sample volume, $V$, calculated:

$$V = 4/3\, p w_{\mathrm{xy}}^2 w_{\mathrm{z}} \qquad\qquad w_{\mathrm{z}} = w\, w_{\mathrm{xy}} \tag{5}$$

The hydrodynamic radius, $R_{\mathrm{H}}$, can be obtained through the Stokes-Einstein equation:

$$R_{\mathrm{H}} = \frac{k_{\mathrm{B}}T}{6phD} \tag{6}$$

where $k_{\mathrm{B}}$ is Boltzmann's constant, $T$ the temperature and $\eta$ the dynamic viscosity of the solvent.

*2.4. Estimation of the number of molecules from the burst photon count distribution*

Photon count distributions were analysed using the model of Fries *et al.* (1), which describes the probability $P(C_B, N_B)$ that a burst contains $C_B$ detected photons, accounting for stochastic emission and molecular dwell times. The mean number of molecules ($N_B$) per population was obtained by fitting this theoretical distribution to the weighted experimental histograms. This analysis yielded independent estimates of the number of molecules for monomeric and aggregated species, respectively.

*2.5. Aggregation analysis*

The aggregation analysis was performed under the theoretical framework developed by our group in previous publications regarding the aggregation of amyloid peptides (2–4). The general equations are shortly displayed here, but the whole theory can be studied in our previous articles.

The total number of protein molecules in the focal volume $NP$ (equivalent to a measure of concentration when divided by the focal volume) is the sum of the numbers of free unlabelled $NP_{\mathrm{f}}^{\circ}$, free labelled $NP_{\mathrm{f}}^{*}$, aggregated unlabelled $NP_{\mathrm{g}}^{\circ}$, and aggregated labelled amyloid $NP_{\mathrm{g}}^{*}$:

$$NP = NP_{\mathrm{f}}^{\circ} + NP_{\mathrm{f}}^{*} + NP_{\mathrm{g}}^{\circ} + NP_{\mathrm{g}}^{*} \tag{7}$$

When we have a mixture of labelled and unlabelled protein in the sample, the labelled protein is randomly distributed among the aggregates, leading to aggregates that bear none, one, or multiple labels. The degree of labelling, $a$, refers to the fraction of fluorescently labelled and unlabelled protein in the sample, which is assumed not to depend on aggregation and follow a Poisson distribution when a monomer occupies an aggregate. Therefore:

$$a = \frac{NP^*}{NP} \tag{8}$$

The mean number of monomers per aggregate (size, or aggregation number) $\bar{n}$ is given by the number of aggregated protein and the number of aggregates, $G$:

$$\bar{n} = \frac{NP_\mathrm{g}}{NG} \tag{9}$$

The fraction of aggregation (degree of aggregation), $\gamma$, refers to the fraction of monomers taking place in an aggregate, and it can be calculated from the labeled fraction alone (assuming the labeling doesn't affect aggregation):

$$\gamma \equiv \frac{NP_\mathrm{g}}{NP} = \frac{NP_\mathrm{g}^*}{NP^*} = \frac{NP^* - NP_\mathrm{f}^*}{NP^*} \tag{10}$$

Knowing $NP^*$, which is the nominal concentration of labelled protein in the sample, it's easy to estimate $NP_\mathrm{f}^*$ from spFCS or the burst photon count distribution and calculate $\gamma$.

When $a$ = 1, which happens when no unlabelled protein is added in the sample, then:

$$\bar{n} = \frac{NP^* - NP_\mathrm{f}^*}{NG^*} \tag{11}$$

$NG^*$ can also be easily obtained from swFCS or the burst photon count distribution.

We can also infer the geometry of the aggregate particles through the FCS and single-molecule data. The diffusion times of homogeneous particles across the FCS sample volume change with their molar mass following a power law, where the exponent $\upsilon$ is related to the geometry of the particle (9, 10). Diffusion times and mean burst duration are also related (1, 7), therefore:

$$t_\mathrm{D} \propto t_\mathrm{B} \propto bM^\upsilon \tag{12}$$

The mean molar mass of the aggregates $M_\mathrm{G}$ is given by the mean aggregation number and the molar mass of the monomers $M_\mathrm{f}$ (disregarding the mass of the labels):

$$M_\mathrm{G} = \bar{n}M_\mathrm{f} \tag{13}$$

The ratio of the diffusion times of aggregates and monomers is then,

$$\frac{t_{\mathrm{B_G}}}{t_{\mathrm{B_f}}} = \frac{b_\mathrm{G}M_\mathrm{G}^{\upsilon_\mathrm{G}}}{b_\mathrm{f}M_\mathrm{f}^{\upsilon_\mathrm{f}}} = \frac{b_\mathrm{G}M_\mathrm{f}^{\upsilon_\mathrm{G}}\bar{n}^{\upsilon_\mathrm{G}}}{b_\mathrm{f}M_\mathrm{f}^{\upsilon_\mathrm{f}}} = \frac{b_\mathrm{G}}{b_\mathrm{f}}M_\mathrm{f}^{(\upsilon_\mathrm{G}-\upsilon_\mathrm{f})}\bar{n}^{\upsilon_\mathrm{G}} \tag{14}$$

$$\log\frac{t_{\mathrm{B_G}}}{t_{\mathrm{B_f}}} = \log\left(\frac{b_\mathrm{G}}{b_\mathrm{f}}M_\mathrm{f}^{(\upsilon_\mathrm{G}-\upsilon_\mathrm{f})}\right) + \upsilon_\mathrm{G}\log\bar{n} \tag{15}$$

The slope of this linear relationship gives information about the geometry of the aggregates.

#### *2.6. Analysis procedure*

The procedure followed to analyse the data from the single-molecule fluorescence experiments was the following:

1. **Define the bursts**. Compute $t_B$ times, apply thresholds, and extract $C_B$, $I_B$, and $I_B$**.**
2. **Population weighting**. Perform Gaussian fits on log histograms of $t_B$and $I_B$; combine into two-dimensional weights.
3. **Weighted FCS**. Compute swFCS curves for each species using photon-weighting.

4. **Model fitting**. Fit correlation curves using Eq. (1), diffusion information from each population is extracted.
5. **Burst distribution fitting**. Apply Fries method (1) and obtain the mean number of molecules through the weighted photon count distribution. $NP_{\mathrm{f}}^{*}$ and $NG^{*}$ are obtained from the monomer and aggregate population, respectively.
6. **Aggregation metrics**. Derive $\gamma$ and $\bar{n}$ from the burst data; estimate aggregate geometry from diffusion scaling.
